# A model for determining the length distribution of actin filaments in cells

**DOI:** 10.1101/2022.03.17.484821

**Authors:** Yuika Ueda, Daiki Matsunaga, Shinji Deguchi

## Abstract

Tensional homeostasis is a cellular process whereby nonmuscle cells such as fibroblasts keep a constant level of intracellular tension and signaling activities. Cells are allowed thanks to tensional homeostasis to adapt to mechanical stress, but the detailed mechanism remains unclear. Here we address from a theoretical point of view what is required for maintaining cellular tensional homeostasis. A constrained optimization problem is formulated to analytically determine the probability function of the length of individual actin filaments (AFs) responsible for sustaining cellular tension. An objective function composed of two entropic quantities measuring the extent of formation and dispersion of AFs within cells is optimized under two constraint functions dictating a constant amount of actin molecules and tension that are arguably the two most salient features of tensional homeostasis. We then derive a specific probability function of AFs that is qualitatively consistent with previous experimental observations, in which short AF populations preferably appear. Regarding the underlying mechanism, our analyses suggest that the constraint for keeping the constant tension level makes long AF populations smaller in number because long AFs have a higher chance to be involved in bearing larger forces. The specific length distribution of AFs is thus required for achieving the constrained objectives, by which individual cells are endowed with the ability to stably maintain a homeostatic tension throughout the cell, thereby potentially allowing cells to locally detect deviation in the tension, keep resulting biological functions, and hence enable subsequent adaptation to mechanical stress. Although minimal essential factors are included given the actual complexity of cells, our approach would provide a theoretical basis for understanding complicated homeostatic and adaptive behavior of the cell.

## 1. Introduction

Homeostasis is responsible for the maintenance of living systems, in which their internal and external elements exhibit complex interactions. In addition to the well-known presence at the organismal level, homeostasis has actually been recognized to be of importance at the individual cellular level. Tensional homeostasis is one of the known homeostatic mechanisms, by which nonmuscle cells such as fibroblasts, endothelial cells, and cancer cells are allowed to sustain a constant level of tension through turnover of their constituent proteins (1–4). Cellular tension is thus relaxed to the basal level over time even upon mechanical stress that disturbs intracellular tensional balance (5, 6). Mechanical stresses such as cell stretch and fluid shear stress are known to trigger signaling pathways, and importantly such signals, typically associated with cell growth and inflammation, are suppressed concomitantly with the recovery of cellular tension. Thus, tensional homeostasis plays crucial roles in functional adaptation of nonmuscle cells to mechanical environment and consequently has been implicated in many diseases such as atherosclerosis (7–12); however, the detailed mechanism remains largely unknown.

Given that living systems are comprised of many structural/functional “layers” that are highly complicated but closely associated with each other, deciphering the homeostasis at the cell layer level will contribute to deeper understanding of the whole homeostasis and resulting adaptive behavior at the organismal level. With this motivation, we are now in the position to tackle this challenging but exciting topic, namely the mechanism underlying cellular tensional homeostasis. Here we discuss physical aspects of the mechanism from a theoretical point of view bridging subcellular and cellular layers. Specifically, we address what is the requirement for actin filaments (AFs), a major cytoskeletal component that undergoes turnover, to allow for sustaining a constant level of tension over the cytoplasm. A constrained optimization problem is then formulated to describe the probability function of the length of individual AFs. Our result yielded a probability function with no local maxima unlike the Gaussian distribution but consistent with experimental observations, describing how the homeostasis at the cellular level is achieved on the underlying basis of the molecular level.

## 2. Method

### 2.1 Constraint function

We aim to derive the length distribution of the individual AFs in cells according to the method of Lagrange multipliers. The number of actin monomers that constitute an AF with a length of *l*_i_ is *n*_i_ where i is a positive integer. We normalize the length (effective diameter) of individual actin monomers to be unity, and AF length is proportional to the number of constituent actin monomers; accordingly, *l*_i+1_ = *l*_i_ + *l*_1_ = *l*_i_ + 1. We consider two equality constraints. First, we assume that the total number of actin monomers within the cell, N, remains constant in accordance with the fact that actin is commonly used as a “housekeeping” protein in Western blotting analysis, and is thus described as

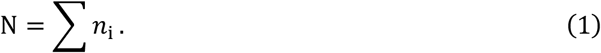

The probability that actin monomers belong to a population of AFs with a length of *l*_i_ is described by

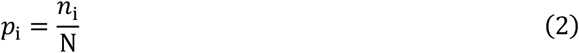

where the sum of the probabilities is unity given Eq. (1).

We impose another equality constraint to capture the essence of tensional homeostasis. Cells are known to sustain preexisting tension of a similar magnitude over the cytoplasm, which is borne by the meshwork of AFs undergoing interactions with myosin II (12–16). These observations suggest that the expected value of the tension in individual AFs of the meshwork is constant. The expected value *F* is then described as an equality constraint by

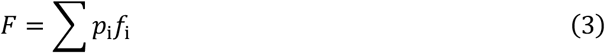

where *f*_i_ is a tension borne by an AF with a length of *l*_i_. The tension in the AF meshwork, which finally gives rise to tensional homeostasis, is generated by the dynamic interaction with myosin II. To bear the tension, individual AFs must be long in length enough to make contact with other surrounding AFs to form a meshwork (17–20). Otherwise, the tension is not transmitted across the cytoplasm even with the presence of myosin II activity. In other words, longer AFs would more frequently bear a level of tension compared to shorter ones. Besides, because there is an upper limit of the myosin II activity, individual AFs would sustain up to a certain level of forces. To capture these features, the relationship between *f*_i_ and *l*_i_ is described by a sigmoid function

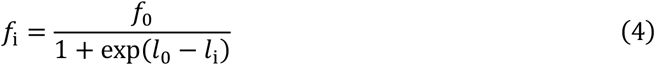

where *f*_0_ and *l*_0_ represent the maximum force and the length with the maximum slope of force, respectively (Supplementary information Fig. S1).

### 2.2 Objective function

What is appropriate for the objective function of the constrained problem that characterizes tensional homeostasis? AFs, responsible for bearing cellular tension, are subject to turnover, by which the constituent monomers are polymerized and/or depolymerized. This continuous turnover allows intracellular AFs to fluctuate in length, and from an entropic perspective they are supposed to take as many states as possible at equilibrium. Thus, the probability distribution of AF length would be partly determined to increase a specific entropy that dictates formation of AFs with extensive fluctuations and is here described by

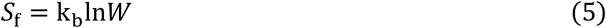

where k_b_ and *W* denote the Boltzmann constant and the number of states, respectively. Under the principle of equal a priori probabilities of each AF length, the number of states of actin molecules is described by

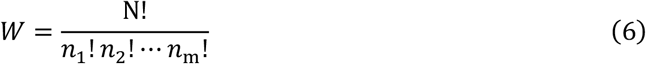

where m is a group of AFs with the longest length. Using Stirling’s approximation and Eqs. (1) and (2), Eq. (5) is rewritten as

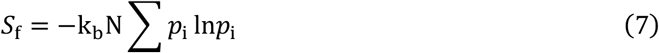

(see Supplementary information for derivation). We adopt this formation-associated entropy as one of the objective functions.

As already discussed above, AFs must be distributed throughout the cytoplasm to form a meshwork where preexisting tension is sustained. As another objective function, we then consider an entropy related to the spatial distribution of AFs given that extensive dispersion of the AFs would allow the cells to be stabilized. Actin-based structures include directional architectures such as stress fibers that align in a specific direction, but the cytoplasmic “directionless” AF meshwork is more commonly present in nonmuscle cells (17–20). The presence of the random meshwork is supposed to allow the cells to stabilize the whole cell architecture (13, 14) as well as to sense and respond to mechanical cues arising from arbitrary directions (11, 21). To evaluate this dispersion-associated entropy, we analyze the extent of intracellular regions where AFs are dispersed depending on their length; here, if all actin monomers form a single long AF, the region of the cell to be occupied, mechanically supported, and communicated by the AF will be limited in space; meanwhile, if actin monomers instead form many short AFs, they overall would be able to cover a large region upon dispersion.

For the analysis, the cytoplasm is modeled to consist of compartments, the number of which is sufficiently larger than that of AFs (Fig. 1). For each AF, the degree of intracellular dispersion is evaluated by counting the maximum number of compartments to which the AF can be distributed. The number of the compartments that an AF can occupy is partly determined by its length *l*_i_ and the compartment size a. We do not consider deformation of AFs for simplicity, and thus only one-dimensional arrangement is analyzed. The maximum number of compartments that can be covered by all the existing AFs is expressed by the following equation (see Supplementary information for detailed explanation):

**Figure 1.**
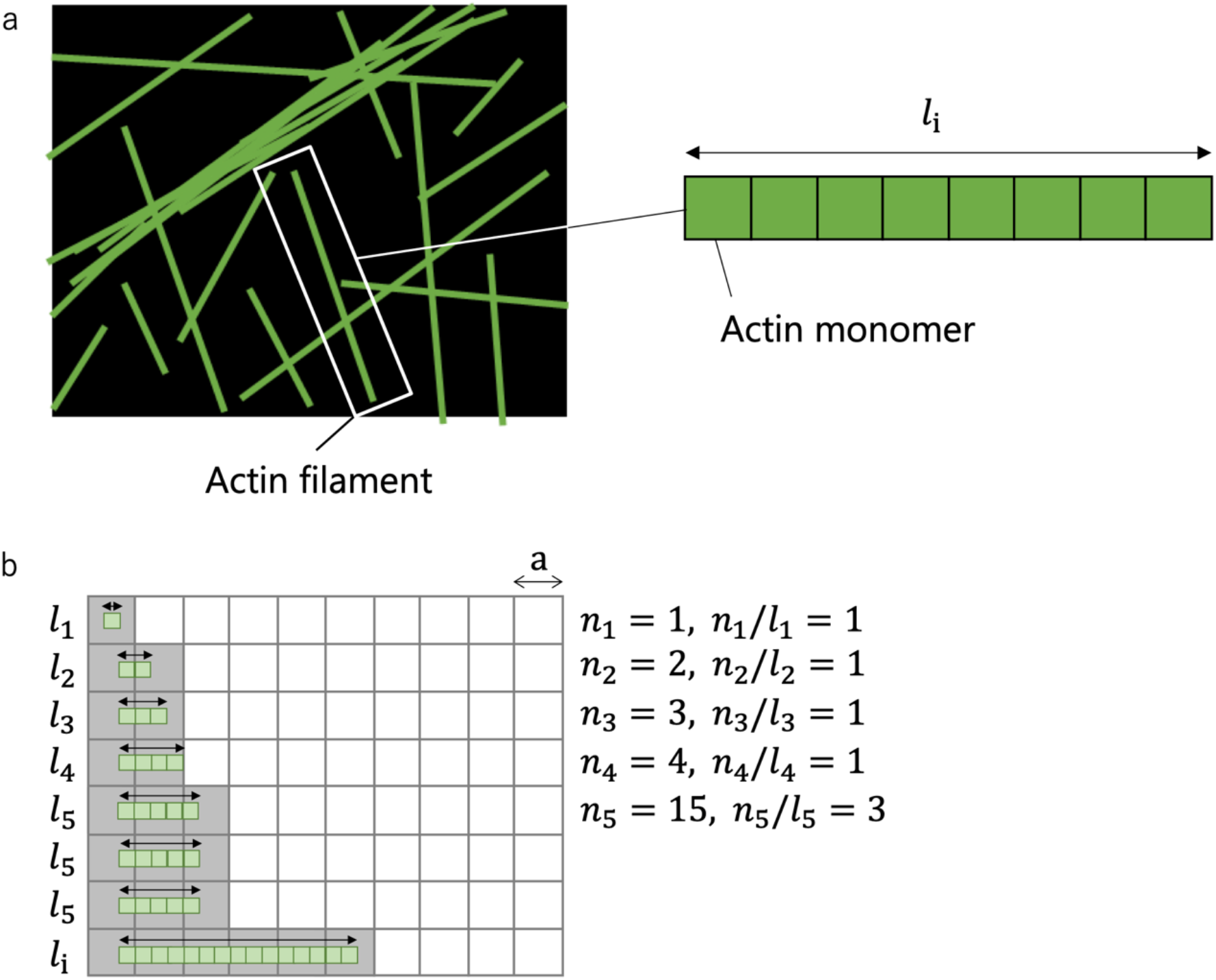
Schematic to explain *S*_*d*_. (a) Schematic of the cytoplasm (black) where AFs (green) assumed to be formed by one-dimensional association of actin monomers to have a length of *l*_i_ are spatially distributed. (b) The cytoplasm is divided by compartments with a length of a. AFs (green) occupy some of the compartments (gray). The arrows in the cell show the length of AFs. An example of *l*_i_ comprising of 15 actin monomers is shown. See Supplementary information for detailed explanation.

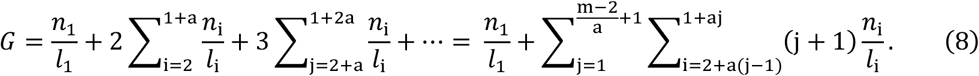

As *n*_i_/*l*_i_ denotes the number of individual AFs of length *l*_i_, we can compute using Eq. (8) the contribution of each length population to *G*, which we denote as *A*_i_. In other words, Eq. (8) is alternatively described as

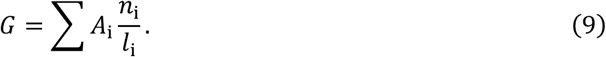

*G* measures the extent of intracellular dispersion of AFs, which is identical in concept to entropy. To evaluate its effect on the stabilization of the cell, we introduce a constant 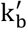 having the same unit as that of the Boltzmann constant k_b_ to define dispersion entropy

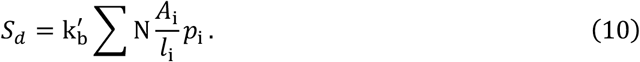

Given that formation entropy *S*_f_ and dispersion entropy *S*_d_ are independent of each other, we describe the overall objective function or the overall entropy of the system *S* as the sum of them, i.e.,

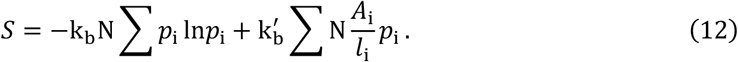

Note that to be exact the overall entropy characterizing highly complex systems like cells would contain numerous entropy-associated factors with a form of the product/sum of them. For example, the product of two hierarchical entropies may be needed to appropriately characterize the overall system if we look at the details of the larger factor. The entropy in such a complex system will thus be practically determined by the final coarseness of the model. The sum, on the other hand, describes independent factors; but, if the effect is small enough, its contribution would be ignored. In the present case, we have considered the two independent entropies, i.e., how AFs are creased (*S*_f_) and how AFs are dispersed (*S*_d_), both of which are supposed be the two most essential factors in the AF-based cellular tensional homeostasis; and, their relative contributions are evaluated by changing the ratio of effective Boltzmann constants k_b_ and 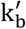, each of which relates the number of their respective possible microstates to the macroscopic cell state.

### 2.3 Lagrange multipliers

Introducing Lagrange multipliers α and β for the constraint function Eq. (1) (divided by N and substituted to Eq. (2)) and Eq. (3), respectively, the partial derivative of the Lagrangian function

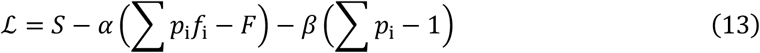

is taken with respect to *p*_*i*_ to find a specific distribution of AFs that maximizes the objective function of Eq. (12) under the constraints. Thus, the probability function of AFs that stabilizes the cell system at equilibrium is obtained by the method of Lagrange multipliers together with thermodynamic considerations to be

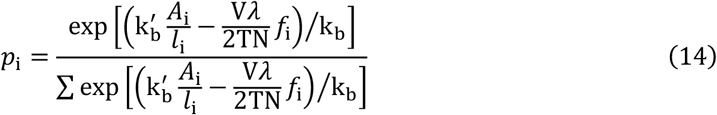

where V, *λ*, and T denote the volume, preexisting strain, and temperature of the cell, respectively (see Supplementary information for derivation).

### 2.4 Parameters

The following parameters are used unless otherwise stated: N = 1,000, k_b_ = 1.38×10^−23^ J/K, T = 309.5 K, V = 6×10^−16^ m^3^, λ = 0.2, *l*_1_ = 1, a = 10, *f*_0_ = 10^−12^ N, and *l*_0_ = 80. Here, the values for T and V were chosen from a typical temperature and cell volume, respectively; the value for N was chosen arbitrarily; the values for the spatial parameters a and *l*_0_ are arbitrarily chosen and are normalized by *l*_1_; the value for k_b_ is taken from the Boltzmann constant; and, the values for λ and *f*_0_ are chosen from in vivo (12–15) and in vitro data (22).

## 3. Results

### 3.1 The relative effect of the two entropies

To capture the basic feature of the appearance probability of actin molecules *p*_i_ derived as a function of length *l*_i_, the effect of changing the ratio of 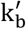 to the constant k_b_ was analyzed. At 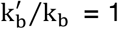, the distribution was approximately comparable in number for all AF lengths (Fig. 2a). As 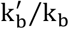 is raised up to 10, the distribution is skewed toward short AFs (Fig. 2b– f). Further smaller and larger 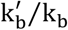 resulted in a distribution similar to that at 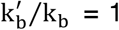 and 10, respectively (Supplementary information Fig. S2). Thus, lowering the relative contribution of 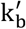, or namely weighting the effect of *S*_f_, gives rise to a uniform distribution of AF length. Meanwhile, increasing the relative contribution of 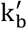, or weighting the effect of *S*_d_, gives rise to a distribution skewed to short AFs. Their log-log plots suggest that, with the raised 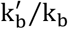, the distribution tends to approach an exponential distribution (Supplementary information Fig. S3). Thus, the overall distribution does not display a peak at a specific length as in a normal distribution and is not preferentially dominated by long AFs.

**Figure 2.**
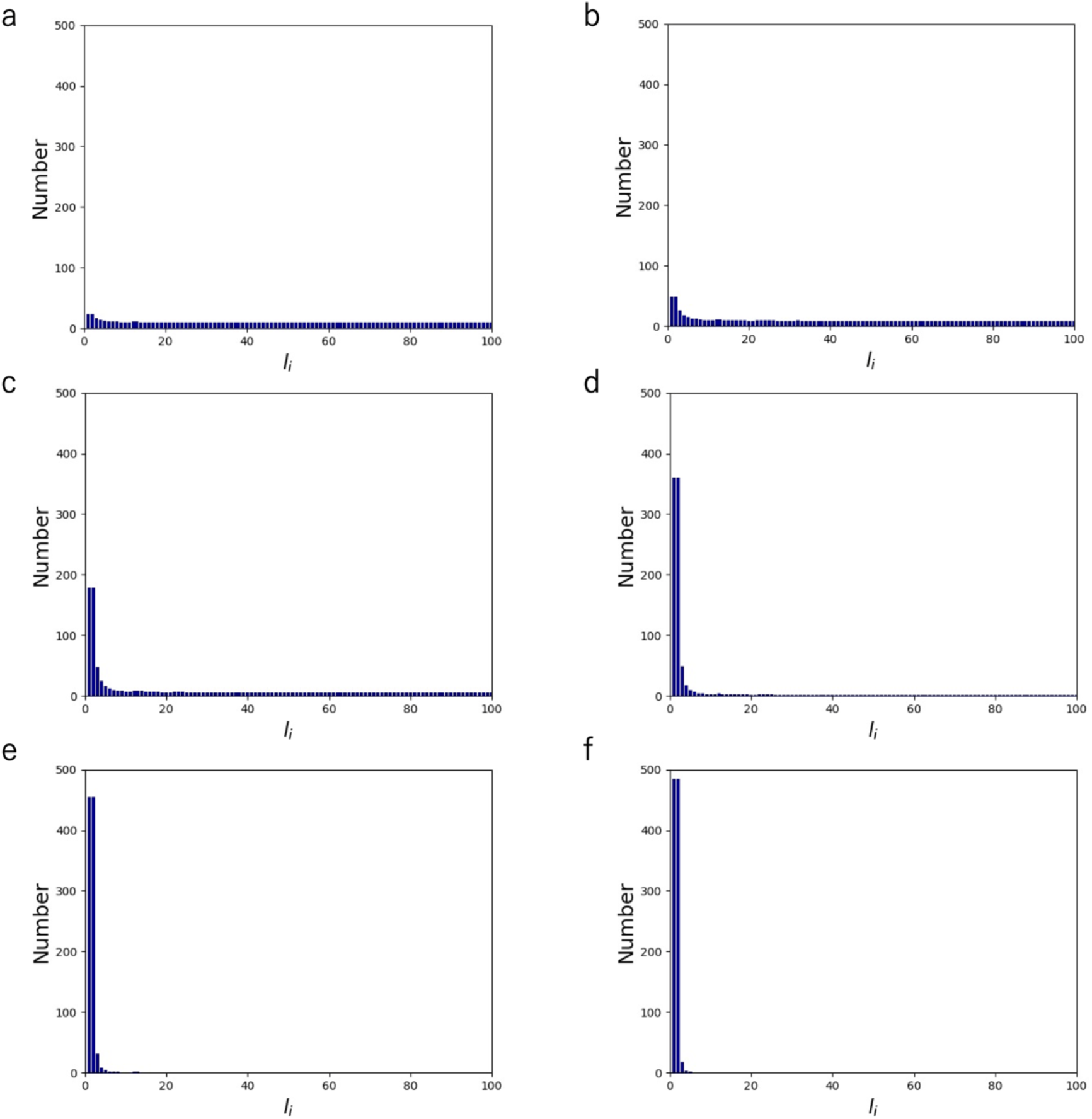
Length distribution of AFs comprising of N = 1,000 monomers at k_b_′/k_b_ = 1 (a), k_b_′/k_b_ = 2 (b), k_b_′/k_b_ = 4 (c), k_b_′/k_b_ = 6 (d), k_b_′/k_b_ = 8 (e), and k_b_′/k_b_ = 10 (f). With a larger k_b_′/k_b_, populations of AFs with shorter lengths increase in the cell.

### 3.2 The effect of force in AFs

The effect of increasing the maximum force *f*_0_ that individual AFs sustain was analyzed at 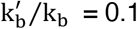 (Fig. 3a) and 4 (Fig. 3b), in which the distribution is uniform and skewed to short AFs, respectively, as already described above. In both cases, the increase in *f*_0_ up to 10^0^ N resulted in suppression of AF populations with a length longer than *l*_0_ (set to be 80 here). To compensate for the suppressed population, the appearance probability was uniformly raised in the other populations. This behavior is observed because of the constraint of the expected value of force given by Eqs. (3) and (4). More specifically, long AFs have a higher chance to be involved in bearing larger forces, and thus the constraint makes long AF populations smaller in number. We also analyzed the effect of changing the threshold length *l*_0_ in Eq. (4) while keeping *f*_0_ unchanged at 10^−12^ N, but almost no change was observed with *l*_0_ = 20 and 80 both at 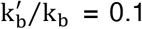 and 4 (Supplementary information Fig. S4). In relation to investigating the role of force, the effect of changing the strain λ, which was introduced in the process of defining the internal energy of the cell from a thermodynamic perspective (see Supplementary information), was also analyzed. The distribution was almost insensitive to the value of λ (0.2 or 0.8) as analyzed at 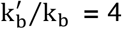 (Supplementary information Fig. S5). The absence of response to these parameters is touched on again in the next section.

**Figure 3.**
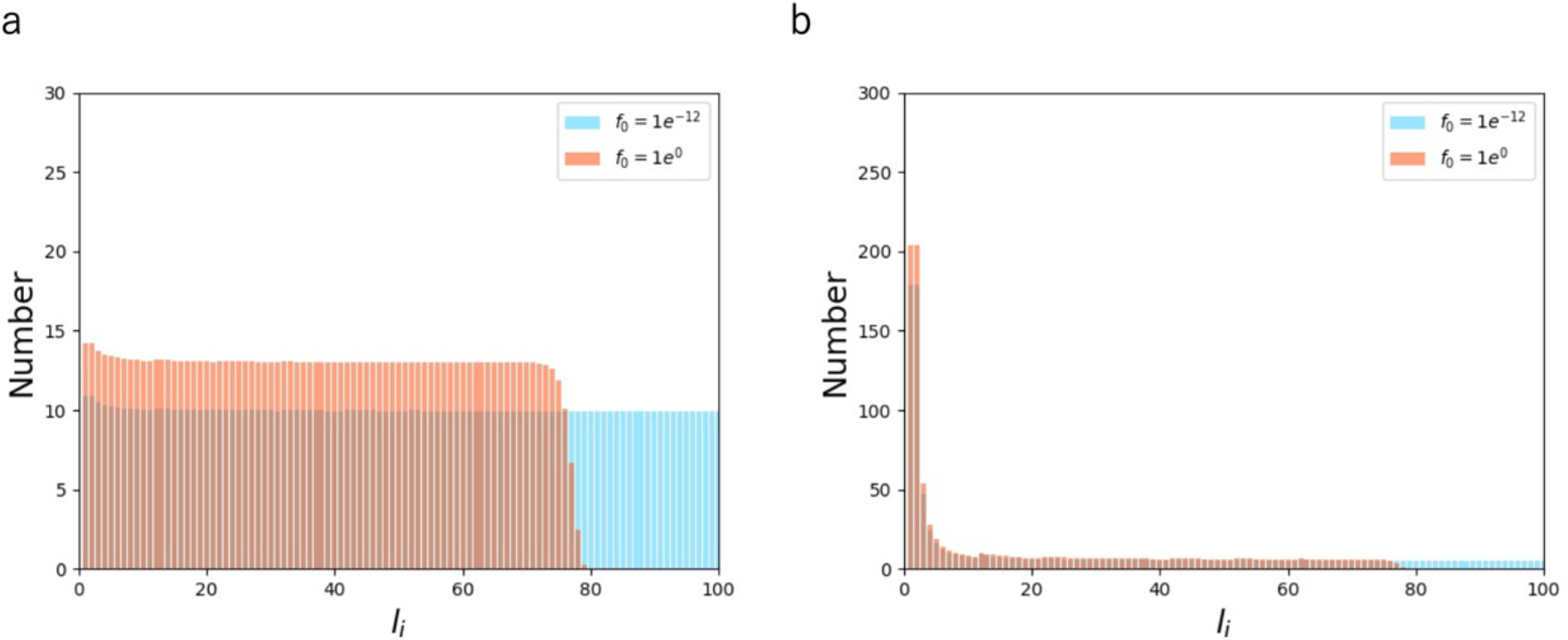
Length distribution of AFs comprising of N = 1,000 monomers with *f*_0_ = 10^−12^ (blue) and *f*_0_ = 1 (orange) at k_b_′/k_b_ = 0.1 (a) and k_b_′/k_b_ = 4 (b).

### 3.3 The individual effect of formation entropy

We have so far considered both *S*_f_ and *S*_d_ for characterizing the overall objective function Eq. (12). To evaluate the sole effect of *S*_f_, here we put 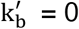 in the probability function of Eq. (14) where the effect of *S*_d_ is omitted. Unlike the case with the two entropies where short AFs appear more frequently (Fig. 2), the distribution becomes uniform all along the length at a maximum force *f*_0_ of more than 10^−4^ N (Fig. 4a, b). As *f*_0_ is increased to 10^−2^ N, a stepwise increase toward shorter AFs appears at the position of *l*_0_ set here to be 80 (Fig. 4c). With a further increase in *f*_0_ to 10^0^ N, the AF population with a length longer than *l*_0_ completely disappears, and accordingly the short population is lifted upward (Fig. 4d). This behavior creating a threshold for appearance is consistent with the case involving the two entropies (Fig. 3) and can thus be explained in the same way as described above; namely, the increase in the maximum force lowers the appearance probability of long AF populations.

**Figure 4.**
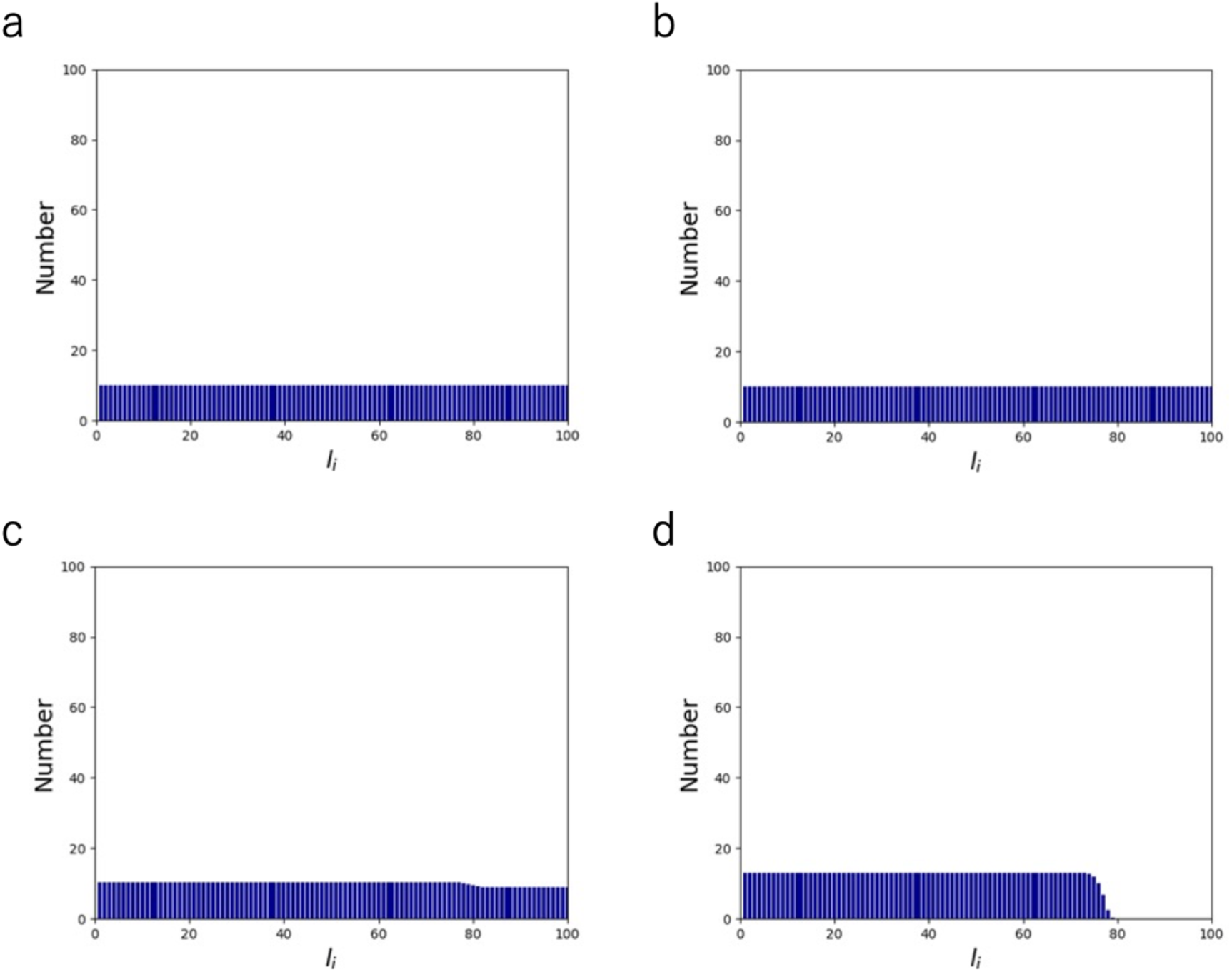
Length distribution of AFs comprising of N = 1,000 monomers at *f*_0_ = 10^−6^ (a), *f*_0_ = 10^−4^ (b), *f*_0_ = 10^−2^ (c), and *f*_0_ = 10^0^ (d).

We next analyzed the effect of changing the threshold length *l*_0_, which had little effect in the presence of *S*_d_ in the objective function (Supplementary information Fig. S4). We found that, in the absence of *S*_d_, only a subtle change is observed at a low *f*_0_ of 10^−4^ N regardless of *l*_0_ = 20 or 80 (Fig. 5a), while the extent of the change becomes profound at an *f*_0_ of 10^0^ N where a steep stepwise change occurs at the position of *l*_0_ (Fig. 5b). Nevertheless, the shape of the uniform distribution does not change except for the position of the threshold for appearance.

**Figure 5.**
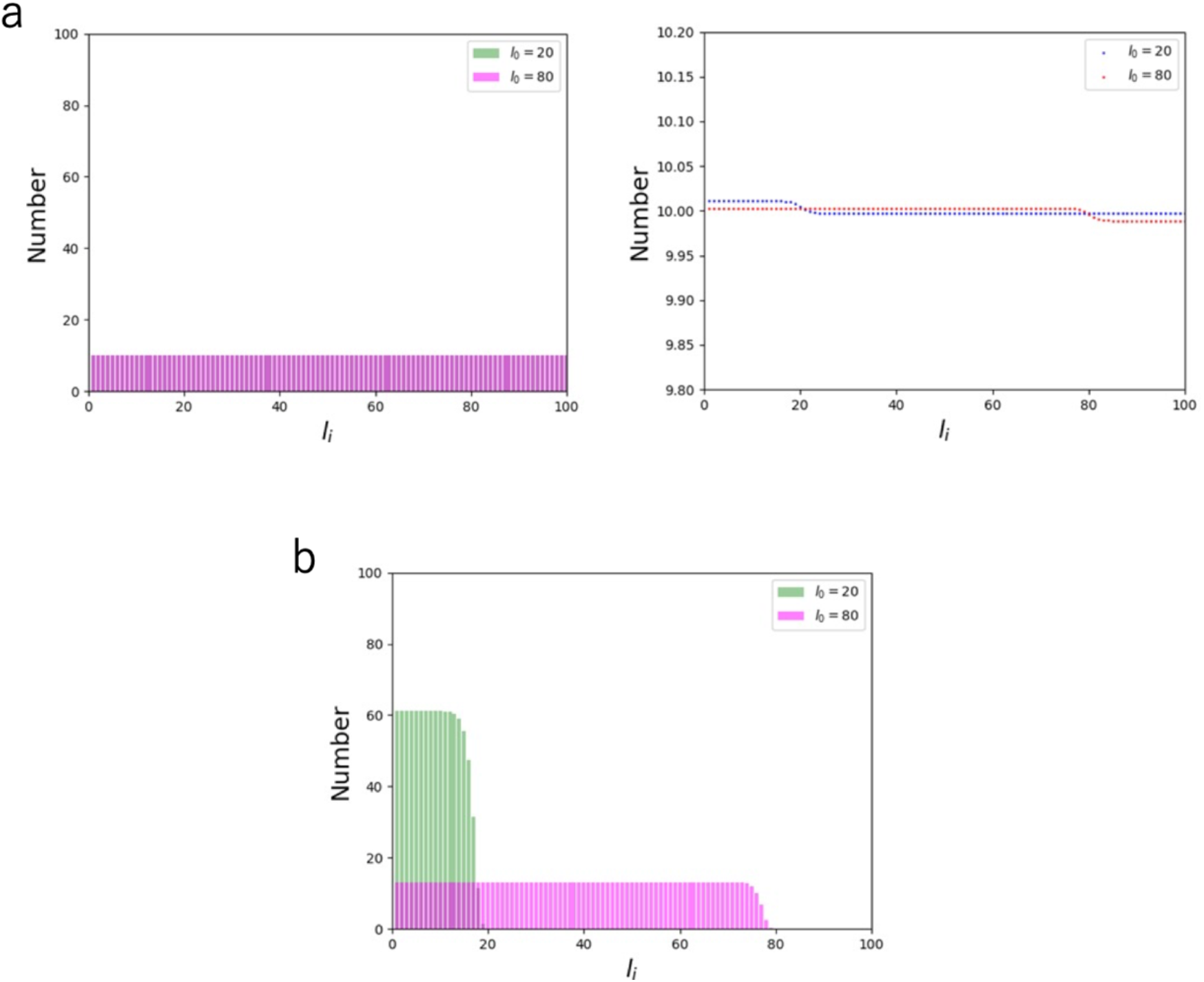
Length distribution of AFs comprising of N = 1,000 monomers with *l*_0_ = 20 (green) and *l*_0_ = 80 (pink) at *f*_0_ = 10^−4^ (a) and *f*_0_ = 10^0^ (b). For visual clarity, point plots are also shown on the right for the case of a with magnified vertical axis.

In relation to investigating the role of force, we analyzed the effect of λ as it partly determines the sensitivity of entropy to force, in which an increased λ is supposed to increase the sensitivity (Supplementary information Eq. S28). Unlike the case with the two entropies where λ has only little effect (Supplementary information Fig. S5), increasing λ from 0.2 to 0.8 does provide a uniform increase in the probability for AF populations with a length shorter than *l*_0_ (Supplementary information Fig. S6). As the increase in λ strengthens the sensitivity to force, it also causes an effect similar to that caused by the increase in *f*_0_, i.e., the uniform lift in probability below *l*_0_ due to the constraint of constant expected value of force (Figs. 3, 4, and 5). The lift becomes particularly detectable in the absence of *S*_d_ as the probability is low and flat over lengths less than *l*_0_, while it is hard to detect the subtle change in the presence of *S*_d_ where the probability is high at short lengths (Fig. S5).

### 3.4 The individual effect of dispersion entropy

Unlike the above case that *S*_d_ is eliminated with 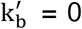, *p*_i_ is not obtained for the case eliminating *S*_f_ as k_b_ is included in the denominator of Eq. (14). As *S*_d_ takes a larger value for shorter AF lengths in a monotonically varying manner (Eq. (10)), the absence of local extrema in the case considering *S*_f_ as the only objective function leads to that the probability is not analytically derived by the method of Lagrange multipliers. The probability that maximizes *S*_f_ under the two constraints dictating the constant amount of actin monomers and force would be one in which shorter AFs more frequently appear. Indeed, increasing the ratio of 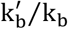 (i.e., decreasing the relative impact of k_b_ on 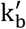) approaches such a distribution (Fig. 2).

In relation to *S*_d_, the effect of *A*_i_ denoting the coefficient of *n*_i_/*l*_i_ in Eq. (9) and measuring how each length population is spatially dispersed was analyzed. The relationship between *A*_i_ and *l*_i_ as a function of the compartment size a shows that a smaller value of a, which more finely divides the cell, increases more steeply with *l*_i_ (Supplementary information Fig. S7a). At a small a, the number of compartments occupied by AFs increases relatively sensitively with increase in AF length; meanwhile, it does not necessarily increase at a large a until the AF length exceeds a certain level to finally make incremental changes. The length required for the stepwise increase is determined by the number of summation operations for i in Eq. (8), specifically (1 + aj) − {2 + a(j − 1)} + 1 = a ; thus, each increment occurs upon a length of a. To evaluate the contribution of each actin molecule to the dispersion with different AF lengths, the number of occupied compartments per unit length (per bound actin molecule), i.e., *A*_i_/*l*_i_, was analyzed (Supplementary information Fig. S7b). *A*_i_/*l*_i_ is large for short AFs and decreases with increased AF length to converge to the reciprocal of a, indicating that the efficiency for intracellular dispersion of individual actin monomers is lowered by being associated with long AFs, but there is a limitation value determined by the compartment size a or equivalently the monomer size. We also analyzed how different values of a affect the length distribution of AFs at different 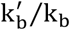 (Figs. S8 and S9). The results show that increasing a as well as 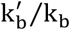 both shifts the distribution toward shorter AF length populations because the increase in a and 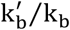 both decreases the efficiency for dispersion (Supplementary information Fig. S7) and the weight on *S*_f_ that leads to a uniform distribution (Fig. 2), respectively. It is important to note that our aim here is to capture the qualitative features of the probability distribution rather than the specific number; thus, as the trend is consistent regardless of the value of a as examined here, we kept a = 10 for the other parts of this study.

## 4. Discussion

Nonmuscle cells such as endothelial cells are known to adapt to mechanical stress applied from arbitrary directions (8, 11, 21). What makes this mechanical adaptation possible is the presence of tensional homeostasis, which allows the cells to sustain a constant level of tension over the cytoplasm through turnover of their constituent proteins (23–25). To address the physical requirement for tensional homeostasis, here we considered what type of actin molecule populations is required to provide a specific mean tension throughout the cytoplasm. The nature of having various AF lengths in nonmuscle cells – which are supposed to behave adaptively in accordance with time-varying intracellular and extracellular milieu – is distinct from that of muscle cells – which are in contrast specialized to exhibit unidirectional contractile activity. Accordingly, for example, nebulin is expressed specifically within muscle cells to stabilize AFs to possess a fixed length (26).

Physicochemical mechanisms determining the length of AFs have been analyzed mostly on their in vitro dynamic behavior where typically the rate of polymerization/depolymerization, viscosity, and diffusion as a function of the length were considered (27–30). AFs in vitro were then predicted to have an exponential distribution, which is qualitatively consistent with experimental observations (31). The effect of some actin-binding proteins was also analyzed, yielding in some cases a distribution with a local maximum (32–37). Regarding the AF distribution in living cells, on the other hand, its involvement in constructing filopodia/lamellipodia was investigated (38). However, previous studies have not explicitly focused on actin meshwork, namely the basic intracellular architecture (17–20). Thus, to our knowledge, there is no study that discussed how individual cells are endowed with tensional homeostasis thanks to their AF resources. Unlike in vitro studies, there are only limited experimental data available on the AF length distribution within animal cells, apart from that in Dictyostelium discoideum (39); but, observations of cell breakage after phalloidin-mediated AF stabilization suggest that the probability is clearly higher for shorter AFs within cells (40), thus consistent with our prediction.

To extract the physical features of tensional homeostasis, we considered the two independent factors: *S*_f_ and *S*_d_ that are related to the formation of the force-bearing elements and to the spatial distribution of the elements, respectively. *S*_f_ indicates in what manner each AF is likely to be formed. For example, if all the molecules have the same length, the number of states is then unity, meaning that they can only be in a fixed state and are thus unstable as a whole since stochastic fluctuations are not allowed. If actin molecules instead take various lengths, they would be more stabilized by having a larger number of states. *S*_d_, on the other hand, indicates in what manner AFs are likely to be dispersed in the cell. Because these two quantities are both identical in concept to entropy, we introduced 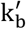 to modulate the weight of *S*_d_ versus *S*_f_, enabling the comparison of the relative impact to the overall system. With regards to the importance of *S*_d_, recent studies have demonstrated that mechanosensitive actin-binding proteins such as filamin are distributed over the cytoplasm (21, 41, 42). These cytoplasmic sensors play a role in maintaining cell functions by converting mechanical information such as intracellular tension into biochemical information such as change in binding affinity, presumably locally detecting deviation in the homeostatic tension and hence keeping the functional integrity of the cell (14). The intracellular dispersion of AFs may thus not only be induced according to the natural law of physics, but also it may be serving for such biological functions.

Studies aimed at characterizing the properties of individual proteins have been extensively conducted as technical advances have made it possible to perform sophisticated measurements at the molecular level, while their collective behaviors are not necessarily taken into sufficient account. However, living organisms are not merely a collection of elements with separate functions but exist through complex interactions among structurally and functionally hierarchical “layers”, thereby allowing the whole system to acquire universal features of life such as flexibility, adaptivity, and resulting maintenance of biological activities. It is necessary, given the actual complexity of cells, to limit the scope of our research to arguably the most essential factor in tensional homeostasis, i.e., the involvement of AFs; but, by describing a simple model, we captured the salient features at the cellular and molecular levels. Specifically, we derived the length distribution of AFs not by considering isolated individuals but by analyzing the population, in which the cellular constraints of having a constant level of actin monomers as well as tension turn out that the probability decreases as the length increases. Our prediction is in agreement with experimental observations made on cells (40). Exploring the involvement of other actin-regulating proteins (43) will be the subject of future challenging but exciting investigations, for which our approach would provide a basis. To this end, as we discussed just after the introduction of the overall entropy of Eq. (12), further elaborate formulations of entropies and appropriate constraints are key to describing the characteristics and roles of the newly added elements.

## Authors’ contributions

Y.U. and S.D. conceived research; Y.U. designed and conducted research with feedback from S.D.; D.M. provided support in analysis; Y.U. and S.D. wrote the manuscript. All authors read and approved the final manuscript.

## Acknowledgments

We thank Tsubasa S. Matsui for discussion.

## Supplementary information

### S1: Figure S1

**Fig. S1.**
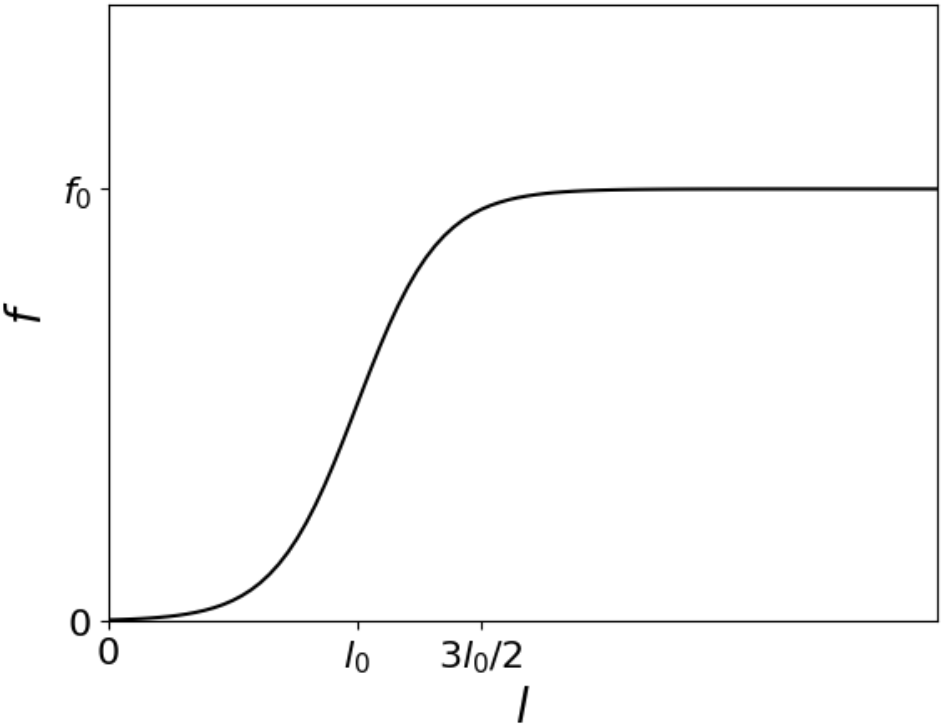
The relationship between force *f*_i_ and AF length *l*_i_. The force increases up to *f*_0_, where *l*_0_ characterizes the position of the transition.

### S2: Derivation of Eq. (7)

From Eqs. (5) and (6), the formation-associated entropy is described by

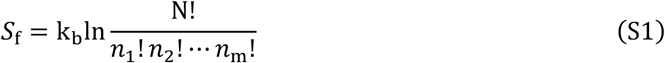

Stirling’s approximation on Eq. (S1) together with Eq. (1) yields that

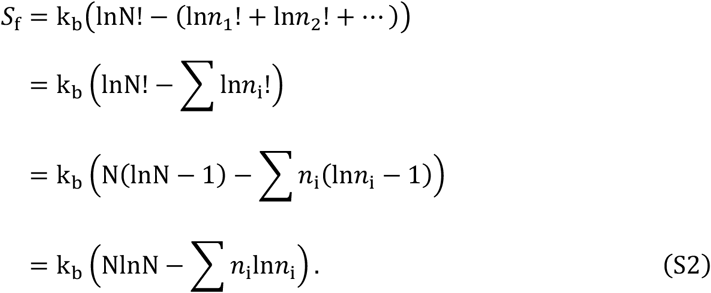

With Eq. (2),

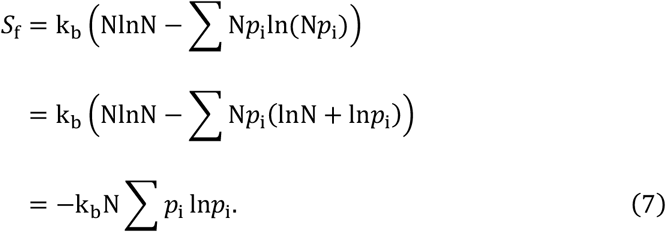

### S3: Explanation of Eq. (8)

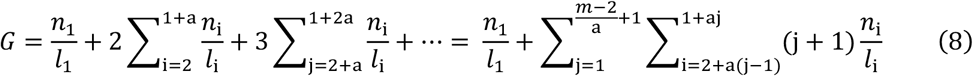

As described in the main text, *n*_i_ is the number of actin monomers in the cell involved in constructing AFs of length *l*_i_. Thus, *n*_i_/*l*_i_ represents the number of individual AFs of length *l*_i_. The diameter of each molecule is smaller than a that is the compartment size, indicating that the number of compartments that can be occupied by a single actin monomer (or an AF of length *l*_1_) is always unity as the monomer is the smallest countable unit, which determines the first term in the middle expression of Eq. (8). Thus, i = 1 represents a group of AFs with the shortest length. As described in the main text, the diameter and filament lengths of actin molecules are normalized by the unit length *l*_1_ = 1. The compartment size of a = 1 means that the diameter of actin molecules is equal to the length of each compartment; thus, the number of compartments occupied by a single AF with a length of, e.g., *l*_i_ = 10 is also 10. Let us consider how the occupied compartments are counted in general with a > 1 based on some examples shown in Fig. 1b where a case with a = 3*l*_1_ = 3 is drawn. Here, the maximum number of compartments that can be occupied by two (*l*_2_), three (*l*_3_), and four (*l*_4_) bound actin molecules is all 2 as indicated by the gray-shaded area in the figure. If actin polymerizes to have a length of *l*_5_ composed of five actin monomers, the maximum number of compartments occupied by this AF is 3 as again indicated by the gray-shaded area. The count for these 2 and 3 consecutive compartments corresponds to the second and third terms in the middle expression of Eq. (8), respectively. Here, the number of the AFs is also considered; namely, in the case of the population of i = 5, the number of the AFs is *n*_5_/*l*_5_ = 15/5 = 3. The general form, in which the maximum number of the compartments occupied by AFs is counted in the same way, corresponds to the right expression of Eq. (8), where m is, again, a group of AFs with the longest length.

**Fig. 1b.**
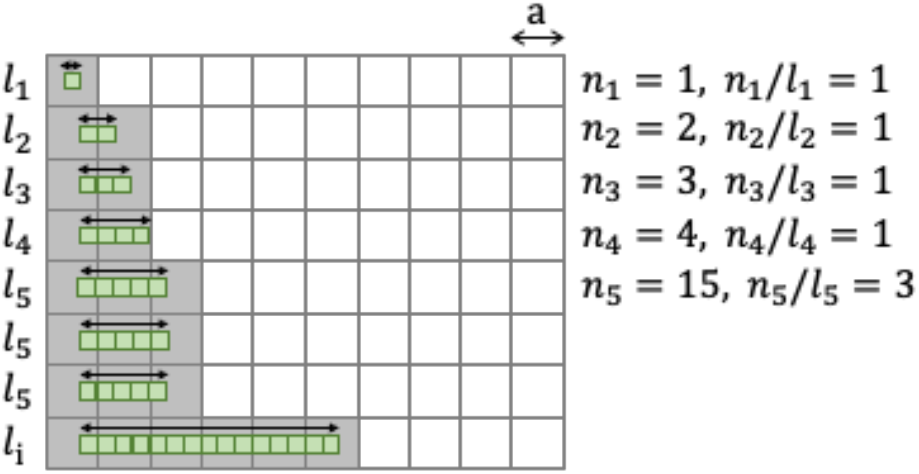

### S4: Derivation of Eq. (14)

The partial derivative of Eq. (13) with respect to *p*_i_ is

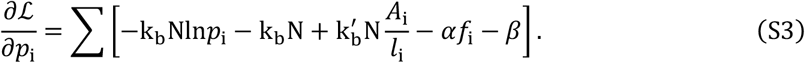

The objective function ℒ takes an extreme value at

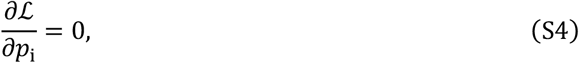

which provides the existence probability for the present purpose

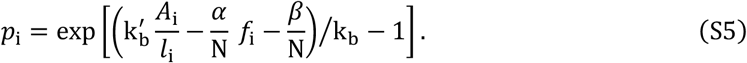

Next, to determine the Lagrange multipliers α and β, Eq. (S5) is substituted partially into Eq. (12) for the overall entropy, yielding

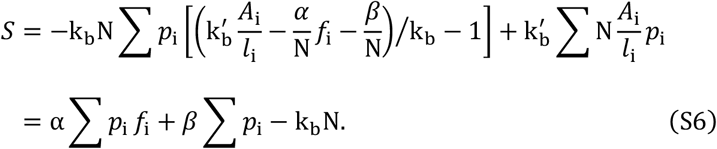

With Eqs. (1)–(3), Eq. (S6) is expressed as

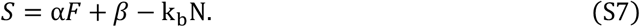

The partial derivative of Eq. (S7) with respect to the expected value of force *F* is

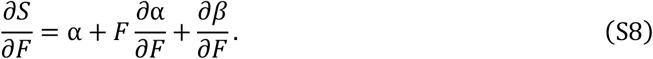

With Eqs. (1) and (2), the summation of *p*_i_ shown by Eq. (S5) must be unity:

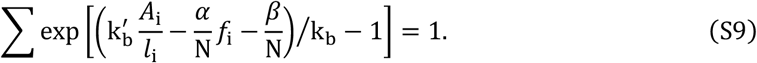

The partial derivative of Eq. (S9) with respect to *F* is

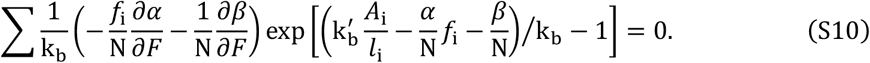

With Eqs. (1), (2), and (S5), Eq. (S10) is reduced to

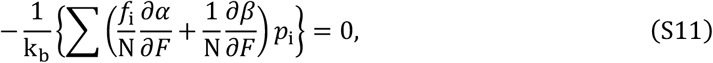

and therefore

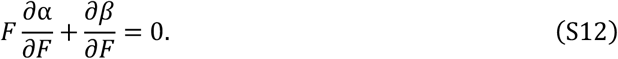

With Eq. (S8), the partial derivative of the overall entropy *S* with respect to *F* is

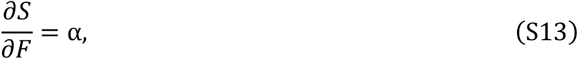

suggesting that the Lagrange multiplier introduced to find the optimal distribution, α, can be expressed using the overall entropy and force expectation value.

Next, we discuss the specific form of α given that the entropy derived from a microscopic point of view based on statistical mechanics, maximizing the objective function at equilibrium, coincides with that derived from a macroscopic point of view based on thermodynamics. For the thermodynamic consideration, we consider a cell with a volume of V and a preexisting strain of λ. As we already discussed, the presence of the preexisting tension (or concomitant mechanical strain) is aimed at capturing the feature of tensional homeostasis. The strain varies spatially and temporarily at the microscopic view, while at the macroscopic view we consider an average level of strain distributed throughout the cytoplasm. The expected value of force *F* introduced in Eq. (3) is regarded as stress in the following thermodynamic model.

The first law of thermodynamics gives

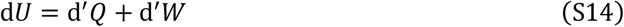

where *U, Q*, and *W* denote the internal energy, the heat given to the system, and the work done to the system, respectively. Let us assume an elastic relationship between stress *F* and strain λ, i.e.,

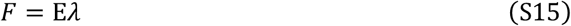

where E denotes the elastic modulus of the cell, and it turns out that the strain energy is

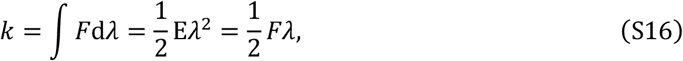

and accordingly the whole strain energy is described using volume V to be

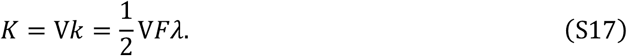

The work caused by change in stress, which in other words is the change in strain energy, is therefore

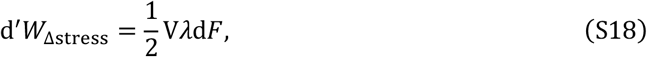

in which cell volume is assumed to be constant (i.e., Poisson’s ratio = 0.5).

The strain or intracellular deformation is generated within cells at tensional homeostasis upon the actin–myosin II interaction using the energy obtained from the ATP-mediated chemical reaction. Therefore, the strain energy discussed here is not the result of work done “by external forces” as in the case of conventional spring elasticity, but is the result of work done “by the system itself.” In other words, as the cell system we discuss here includes subcellular parts that do work (namely, generate force and deform), the work done to the system must be opposite in sign to that done “by the system itself.” It also turns out that, in the present system, the work done “by the system itself” will increase in amount as the strain/deformation-associated intracellular stress *F* increases. Therefore, change in work of the entire system is

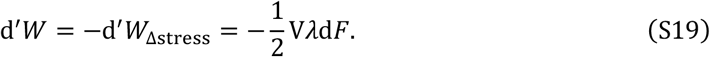

With this, Eq. (S14) turns out

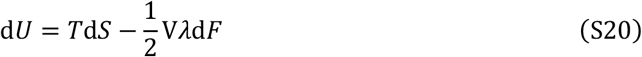

where *T* is the thermodynamic temperature. The free energy *G* is defined as

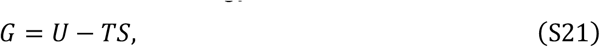

and its change is

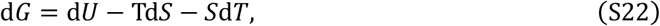

or with Eq. (S20)

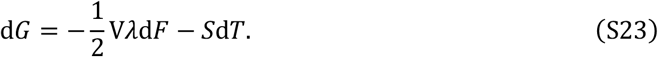

Describing the total differential form of *G*(*F, T*),

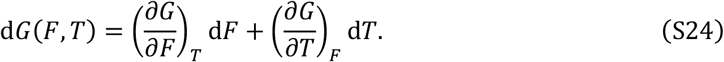

From Eqs. (S23) and (S24),

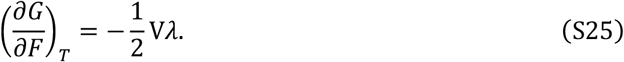

The partial derivative of the free energy with respect to *F* at constant temperature yields

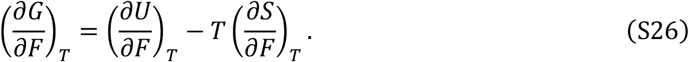

From Eqs. (S25) and (S26),

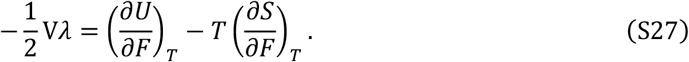

At constant temperature where there is no change in internal energy, Eq. (S27) is reduced to

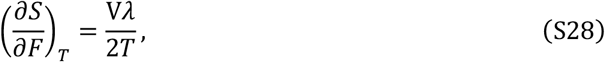

describing the effect of intracellular stress on entropy derived from a macroscopic thermodynamic point of view. This positive relationship – distinct from the negative one for a conventional spring where stretch reduces the extent of fluctuations, the number of possible microstates of the constituents, and thereby the entropy as well – is reasonable because the stress of current interest *F* is, as already mentioned, originated from the intracellular actin– myosin II interaction. More specifically, *F* is interpreted to be an indicator of the activity of the intracellular components rather than a factor compelling their movement. Thus, an increased *F* and resulting activation of the actin–myosin II interaction stabilize the cell structure as well as increase the entropy. Given that Eqs. (S13) and (S28) are equal at equilibrium,

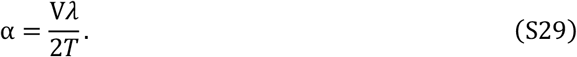

Substituting Eq. (S29) into (S5) yields

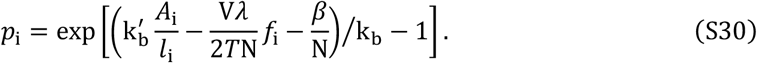

To determine *β*, the constant part of Eq. (S30), which is independent of i, was expressed as B, and then

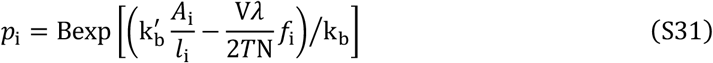

where

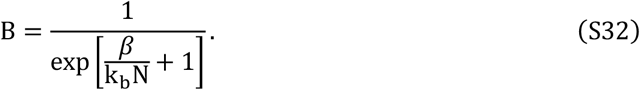

With Eqs. (1) and (2), the summation of *p*_i_ must be unity so that

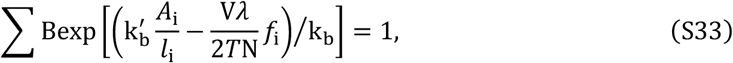

indicating that

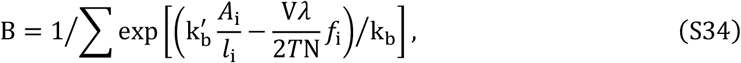

and

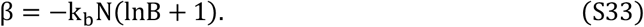

Consequently,

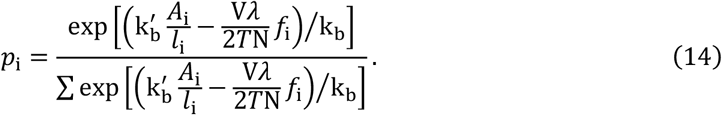

### S5: Figure S2

**Fig. S2.**
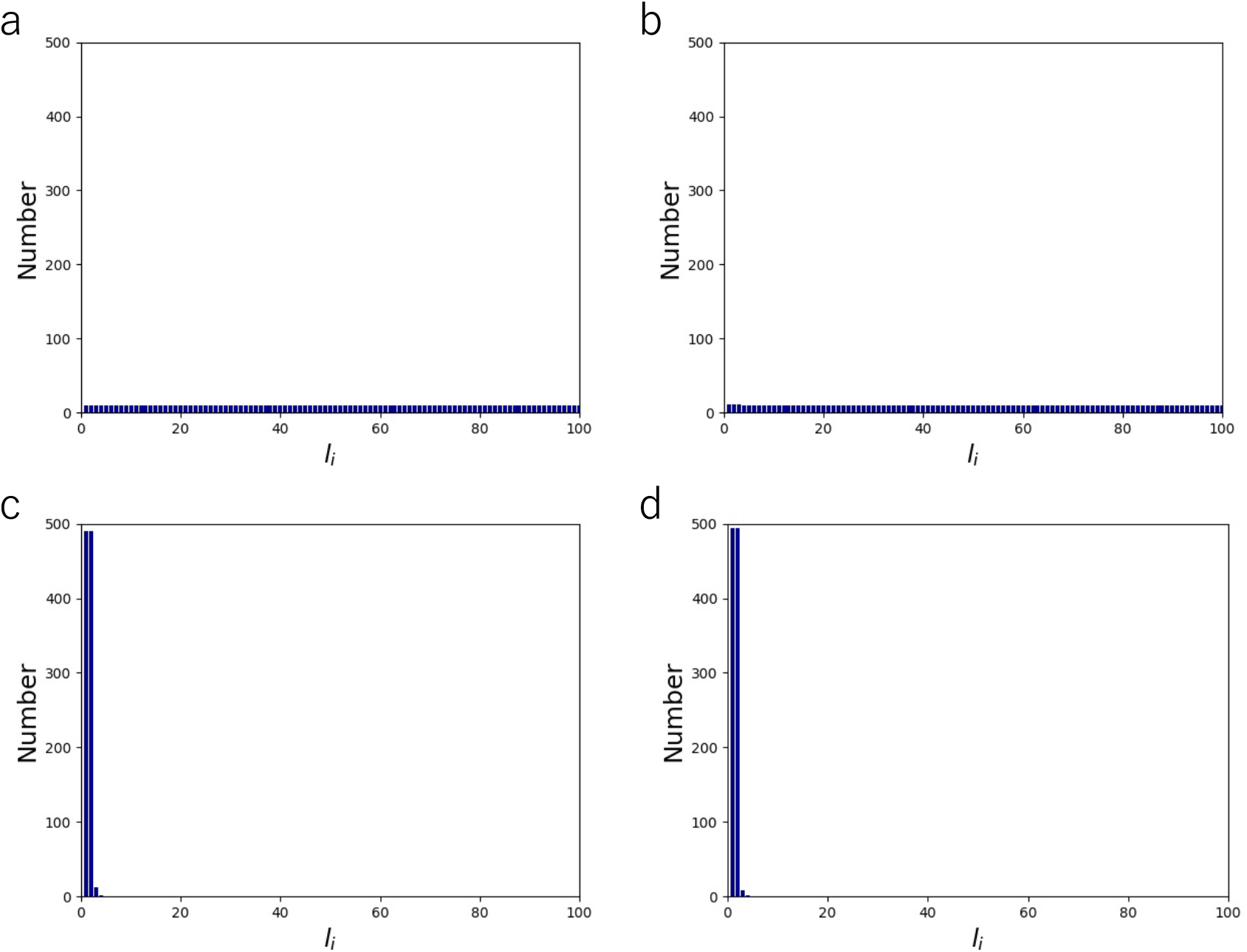
Length distribution of AFs comprising of N = 1,000 monomers at k_b_′/k_b_ = 10^−2^ (a), k_b_′/k_b_ = 10^−1^ (b), k_b_′/k_b_ = 11 (c), and k_b_′/k_b_ = 12 (d).

### S6: Figure S3

**Fig. S3.**
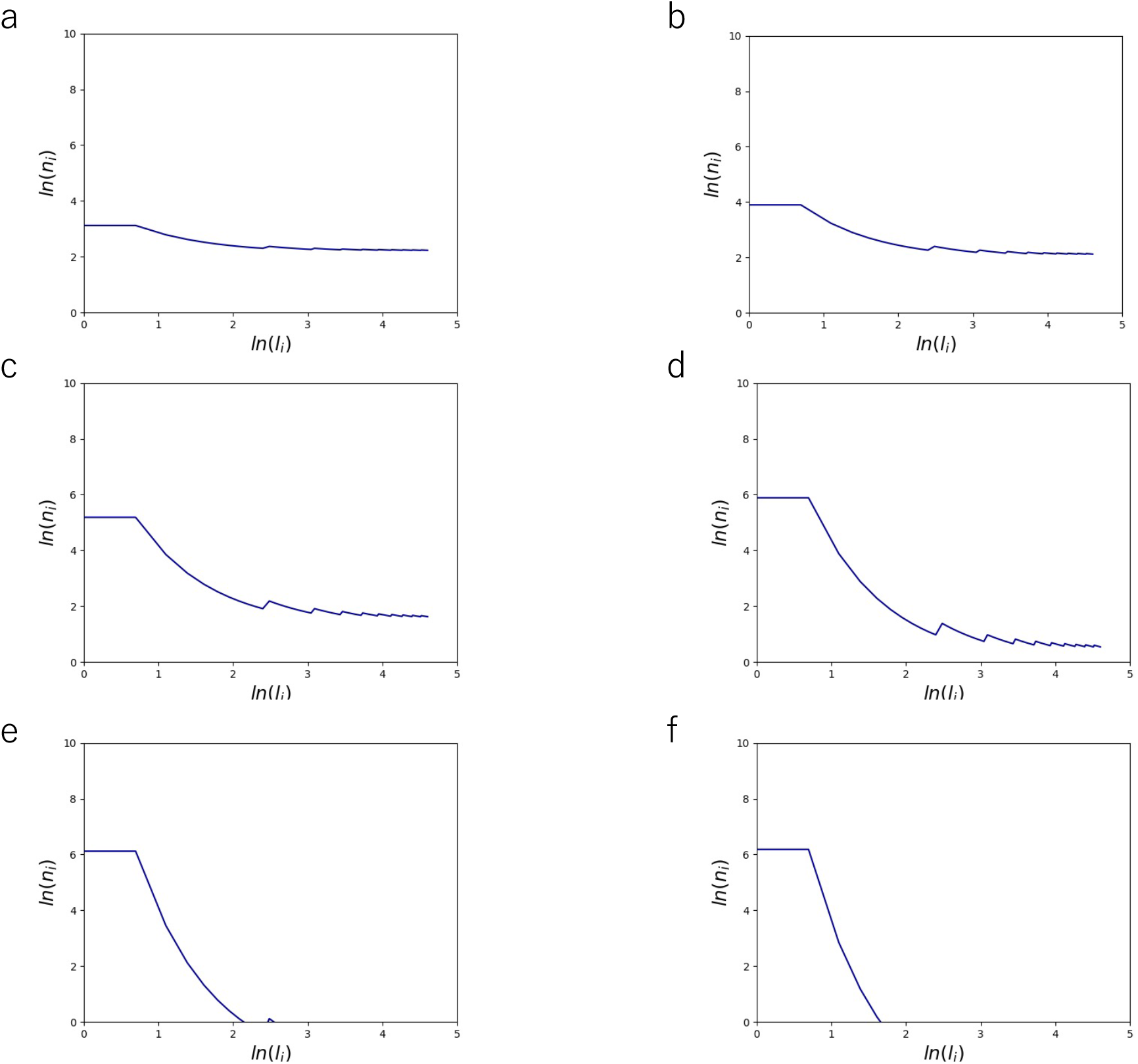
Log-log graphs of Fig. 2, showing the length distribution of AFs comprising of N = 1,000 monomers at k_b_′/k_b_ = 1 (a), k_b_′/k_b_ = 2 (b), k_b_′/k_b_ = 4 (c), k_b_′/k_b_ = 6 (d), k_b_′/k_b_ = 8 (e), and k_b_′/k_b_ = 10 (f).

### S7: Figure S4

**Fig. S4.**
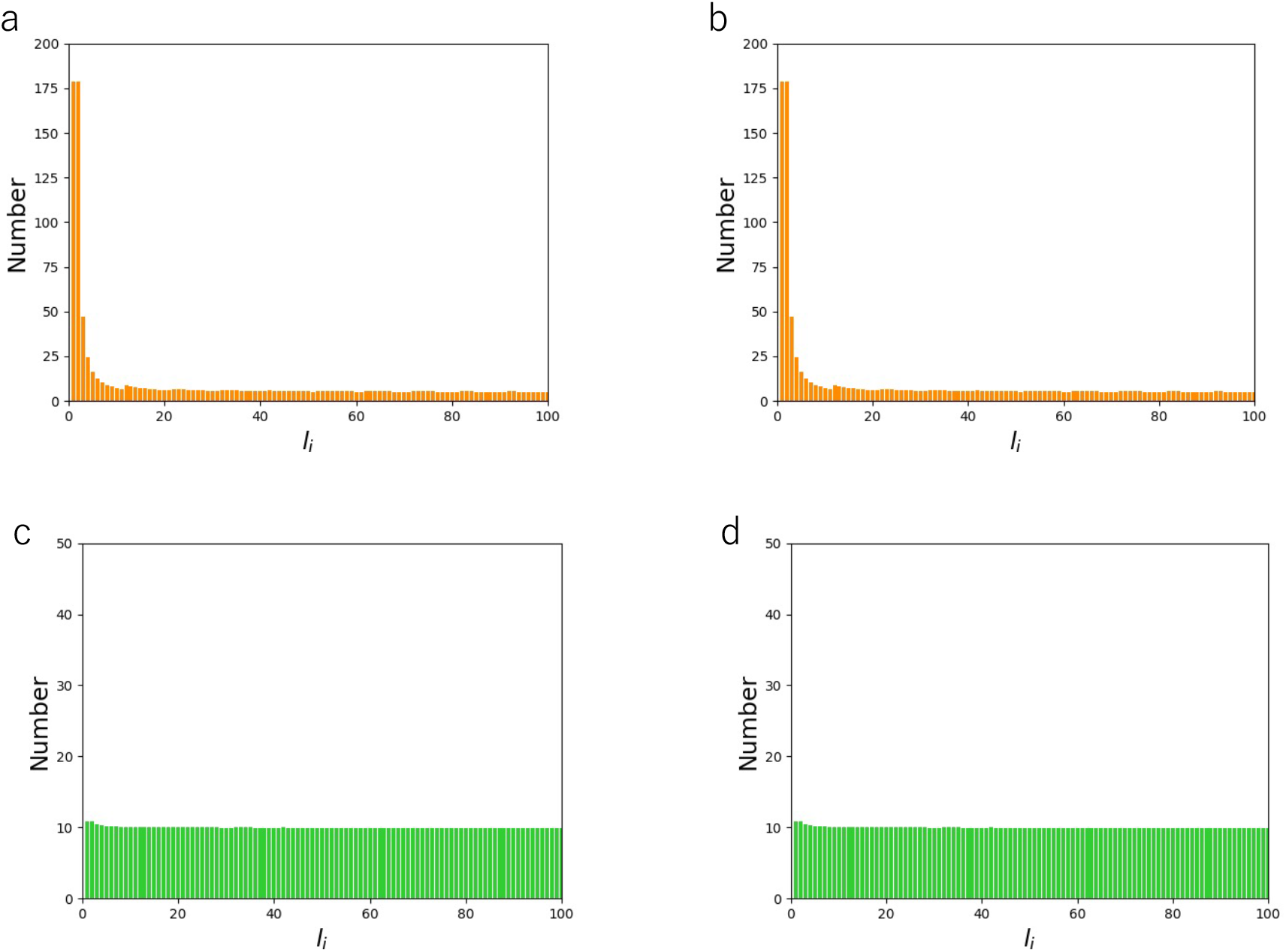
Length distribution of AFs comprising of N = 1,000 monomers at k_b_′/k_b_ = 4 and *l*_0_ = 20 (a), k_b_′/k_b_ = 4 and *l*_0_ = 80 (b), k_b_′/k_b_ = 0.1 and *l*_0_ = 20 (c), and k_b_′/k_b_ = 0.1 and *l*_0_ = 80 (d), showing that the distribution does not depend on *l*_0_.

### S8: Figure S5

**Fig. S5.**
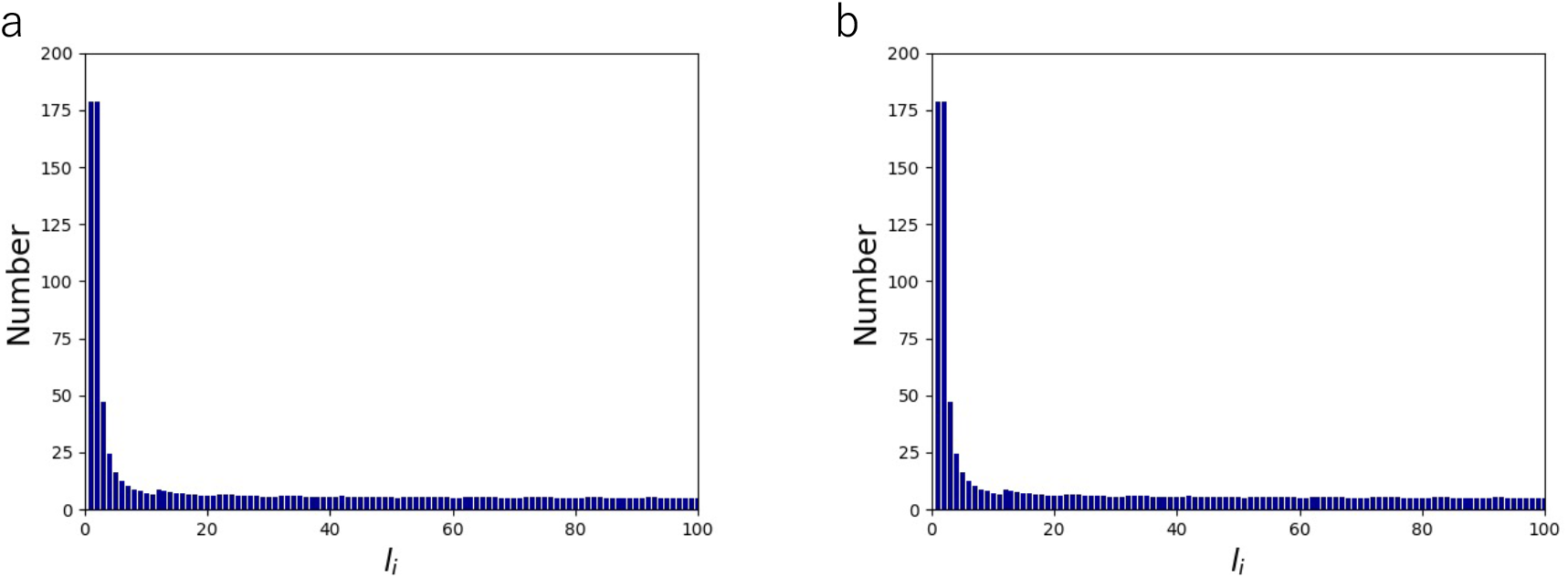
Length distribution of AFs comprising of N = 1,000 monomers at preexisting cell strain (prestretch) of λ = 0.2 (a) and λ = 0.8 (b).

### S9: Figure S6

**Fig. S6.**
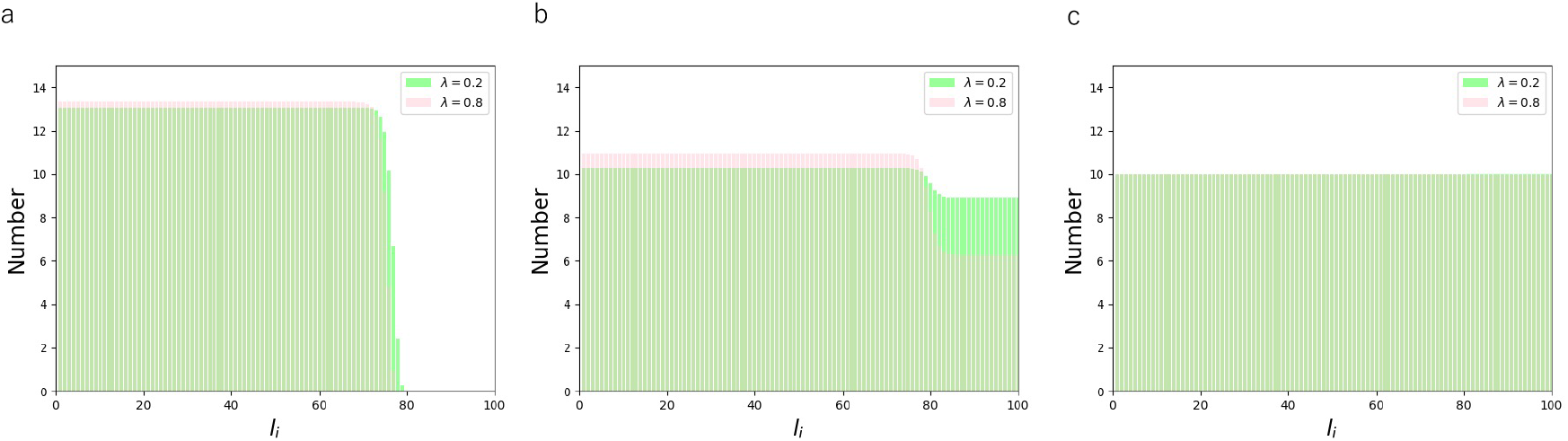
Length distribution of AFs comprising of N = 1,000 monomers for preexisting cell strain of λ = 0.2 (green) and λ = 0.8 (pink) at *f*_0_ = 10^0^ (a), *f*_0_ = 10^−2^ (b), and *f*_0_ = 10^−4^ (c).

### S10: Figure S7

**Fig. S7.**
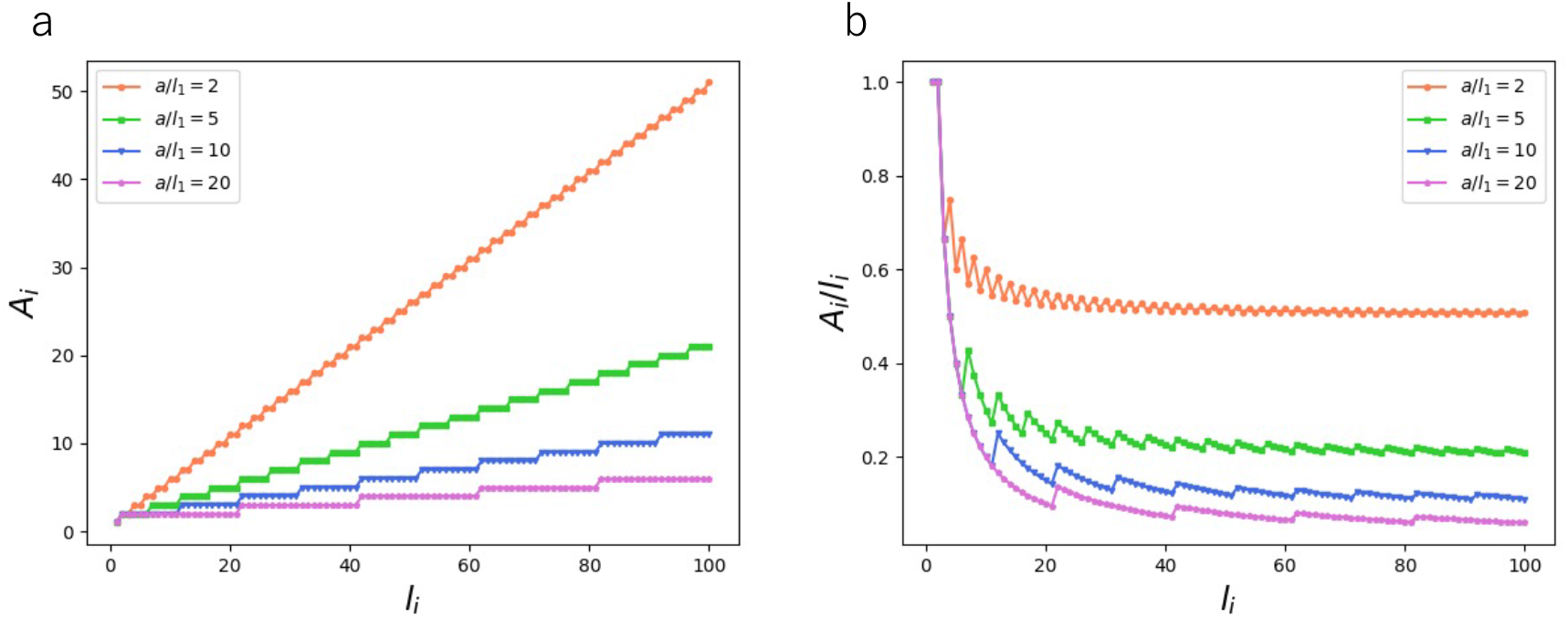
The effect of changing the ratio of compartment size a to actin monomer size *l*_1_. In the analysis, the size is normalized by *l*_1_ as described in Methods section. (a) *A*_i_, which determines *S*_d_, as a function of *l*_1_ at different a; a/*l*_1_ = 2 (orange), a/*l*_1_ = 5 (green), a/*l*_1_ = 10 (blue), and a/*l*_1_ = 20 (purple). (b) *A*_i_/*l*_*i*_ as a function of *l*_1_ at different a; a/*l*_1_ = 2 (orange), a/*l*_1_ = 5 (green), a/*l*_1_ = 10 (blue), and a/*l*_1_ = 20 (purple). Note that *l*_1_ = 1 for the rest of the analyses in the present study.

### S11: Figure S8

**Fig. S8.**
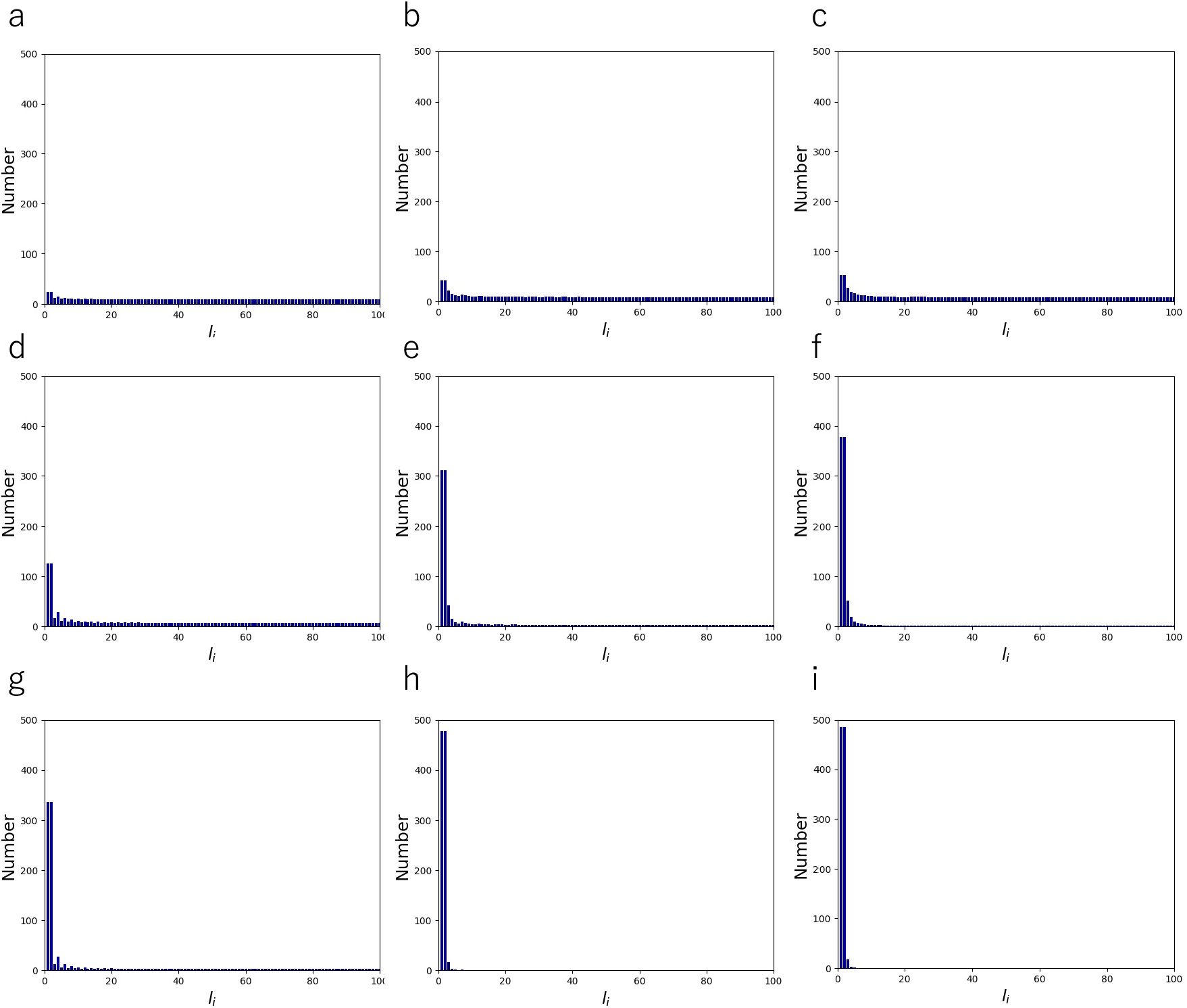
Length distribution of AFs comprising of N = 1,000 monomers at k_b_′/k_b_ = 2 and a/*l*_1_ = 2 (a), k_b_′/k_b_ = 2 and a/*l*_1_ = 5 (b), k_b_′/k_b_ = 2 and a/*l*_1_ = 20 (c), k_b_′/k_b_ = 6 and a/*l*_1_ = 2 (d), k_b_′/k_b_ = 6 and a/*l*_1_ = 5 (e), k_b_′/k_b_ = 6 and a/*l*_1_ = 20 (f), k_b_′/k_b_ = 10 and a/*l*_1_ = 2 (g), k_b_′/k_b_ = 10 and a/*l*_1_ = 5 (h), and k_b_′/k_b_ = 10 and a/*l*_1_ = 20 (i). Cases at a/*l*_1_ = 10 are shown in Fig. 2.

### S12: Figure S9

**Fig. S9.**
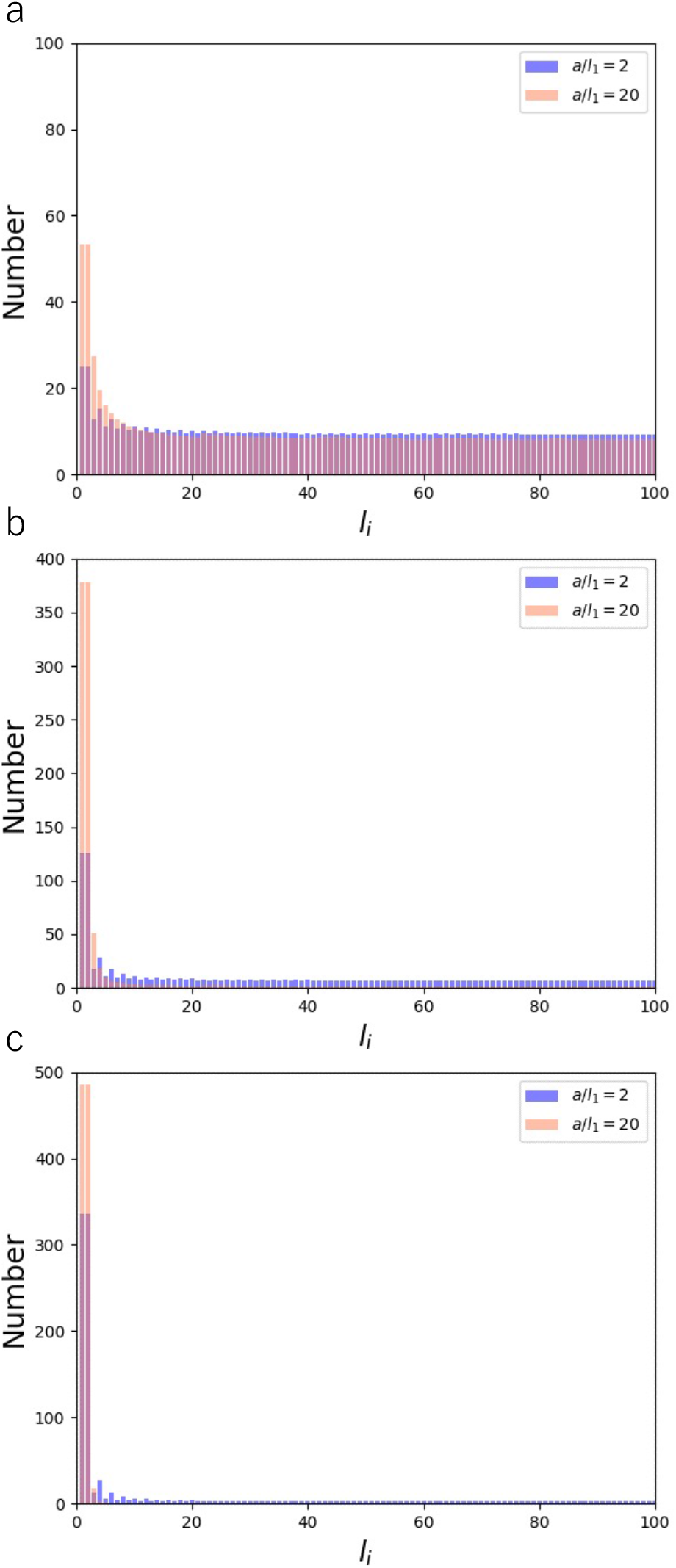
Length distribution of AFs comprising of N = 1,000 monomers for a/*l*_1_ = 2 (blue) and a/*l*_1_ = 20 (orange) at k_b_′/k_b_ = 2 (a), k_b_′/k_b_ = 6 (b), and k_b_′/k_b_ = 10 (c) taken partly from Fig. S8.

